# Disentangling the Role of Face Typicality and Affect in Emotional Face Processing: Self-reported and Electrophysiological Evidence

**DOI:** 10.1101/378224

**Authors:** Mariana R. Pereira, Tiago O. Paiva, Fernando Barbosa, Pedro R. Almeida, Eva C. Martins, Torsten Baldeweg, Michelle de Haan, Fernando Ferreira-Santos

**Author notes:** Correspondence concerning this article should be addressed to Fernando Ferreira-Santos, Laboratory of Neuropsychophysiology, Faculty of Psychology and Education Sciences, University of Porto, R. Alfredo Allen, 4200-135 Porto, Portugal. This research was funded by the BIAL Foundation (Grant 242/14).

## Abstract

Typicality, or averageness, is one of the key features that influences face evaluation, but the role of this property in the perception of facial expressions of emotions is still not fully understood. Typical faces are usually considered more pleasant and trustworthy, and neuroimaging results suggest typicality modulates amygdala and fusiform activation, influencing face perception. At the same time, there is evidence that arousal is a key affective feature that modulates neural reactivity to emotional expressions. In this sense, it remains unclear whether the neural effects of typicality depend on altered perceptions of affect from facial expressions or if the effects of typicality and affect independently modulate face processing. The goal of this work was to dissociate the effects of typicality and affective properties, namely valence and arousal, in electrophysiological responses and self-reported ratings across several facial expressions of emotion. Two ERP components relevant for face processing were measured, the N170 and Vertex Positive Potential (VPP), complemented by subjective ratings of typicality, valence, and arousal, in a sample of 30 healthy young adults (21 female). The results point out to a modulation of the electrophysiological responses by arousal, regardless of the typicality or valence properties of the face. These findings suggest that previous findings of neural responses to typicality may be better explained by accounting for the subjective perception of arousal in facial expressions.

---

The human face carries important information in a social context, allowing us to infer aspects such as the other’s identity and emotional state (Bruce & Young, 1986), and to draw social judgments (Willis & Todorov, 2006). Face typicality is one of the factors that influence face evaluation. A face is more typical the closer it is to a prototypical configuration, which is conceived as an average of all the past-perceived faces (Diamond & Carey, 1986; Valentine, 1991). Therefore, in a face-space each face may be represented as a vector that originates from the central face and the further the face is from the average face, the more atypical it is (Rhodes & Leopold, 2011). Recent work on face perception, based on Predictive Processing approaches, is largely consistent with these models and perhaps complements them with important insights as to the neural basis of these processes (Brodski-Guerniero, Paasch, Wollstadt, Özdemir, Lizier, & Wibral, 2017; Trapp, Schweinberger, Hayward, & Kovács, 2018). From a Predictive Processing perspective, face perception will depend on the comparison between the visual sensory input of the present face stimulus and a prior internal model (prediction) of the relevant object features. The mismatch between the sensory input and the prior will result in a prediction error that is processed as the level of novelty of the stimulus (Arnal & Giraud, 2012). In this view, face typicality could consist of a measure of the difference between a given face stimulus and the individual’s internal model of a general face that would serve as a prior expectation for face perception.

As Sofer and colleagues mention (2015), the typical face carries a special status perceptually and socially. For instance, typical faces are considered more trustworthy, explained by the inherent familiarity of any typical stimuli, associated with positive affect (Lee, 2001) and trustworthiness (Faerber, Kaufmann, Leder, Martin, & Schweinberger, 2016). Face typicality has also been associated with attractiveness, although the results are incongruent, with studies finding typical faces as more attractive (e.g., Langlois & Roggman, 1990; Rhodes, Sumich, & Byatt, 1999; Valentine, Darling, & Donnelly, 2004) whereas others find the opposite effect (e.g., de Bruine, Jones, Unger, Little, & Feinberg, 2007).

Typicality also seems to play a part in the neural mechanisms of face evaluation. Previous studies found that amygdala and fusiform facial area (FFA), regions involved in facial and affective processing (Kanwisher & Yovel, 2006; Todorov, 2012), show a non-linear modulation by both attractiveness (Chatterjee, Thomas, Smith, & Aguirre, 2009; Iaria, Fox, Waite, Aharon, & Barton, 2008; Shen et al.,2016; Winston, O’Doherty, Kilner, Perrett, & Dolan, 2007) and trustworthiness (Said, Baron, & Todorov, 2009; Santos, Almeida, Oliveiros, & Castelo-Branco, 2016). However, Mende-Siedlecki, Said, and Todorov (2013) meta-analysed neuroimaging studies on facial evaluation and suggested that the non-linear results may be explained by differences in the typicality of the faces used in the studies (see also Mende-Siedlecki, Verosky, Turk-Browne, & Todorov, 2013). Indeed, a higher responsivity of the amygdala and FFA was already found to atypical comparably to typical faces, independently of their social features and valence properties (Mattavelli, Andrews, Asghar, Towler, & Young, 2012; Said, Dotsch, & Todorov, 2010; Todorov & Engell, 2008).

However, the affective arousal properties of the stimuli may play a part in this increased activation to atypical faces, and not typicality itself. Indeed, atypical stimuli represent a bigger potential threat than familiar ones, which possibly alters their perceived arousal, increases their relevance, and demands the allocation of more resources (discussed in Mende-Siedlecki, Verosky, Turk-Browne, & Todorov, 2013). Additionally, there is already evidence regarding the role of perceived arousal of various stimuli in the modulation of amygdala activation, regardless of the valence (Kensinger & Schacter, 2006; Sergerie, Chochol, & Armony, 2008; Zald, 2003). These results also are in accordance to previous findings of a U-shaped amygdala activation to both highly pleasant and unpleasant sensory stimuli (Anderson et al 2003; Small et al., 2003), which convey higher levels of activation than neutral stimuli. These results may hold relevance for visual face processing as the amygdala is known to enhance visual cortical processing for expressive faces (Hadj-Bouziane et al., 2012; Vuilleumier, Richardson, Armony, Driver, & Dolan, 2004).

Overall, the roles of typicality and affective properties of facial stimuli have been approached as modulators of brain activity and face evaluation, but their overlap is still undefined. In this study, we aimed to address this issue by dissociating typicality from the affective properties, specifically arousal and valence, and to examine the effect of these factors in the modulation of electroencephalographic activity. To do this, we used a design in which participants were exposed to faces manipulated across three experimental factors: (1) faces could be typical or atypical, (2) portray different emotional categories, and (3) vary in affective properties (arousal or valence) within each category. This design should allow disentangling the effects of typicality from those of affective properties, shedding light on the hemodynamic and electrophysiological findings reviewed above.

Regarding self-reports we were interested in the individual ratings of arousal, valence, and typicality to each stimulus, in order to assess a possible impact of typicality in affective perception, and vice-versa. In terms of neurophysiological indexes, we chose an ERP methodology due to the high temporal resolution of this technique, allowing to understand the neural dynamics locked to the events. The N170 is one of the most studied components in the field, defined as a face-sensitive negative potential that peaks between 150ms and 220ms poststimulus in occipito-parietal sites, bilaterally. This component registers changes in latency and amplitude with inverted and scrambled faces comparably to upright and normative facial configurations (Bentin, Allison, Puce, Pereza, & McCarthy, 1996). These changes may indicate its modulation by differences in the structural configuration of the face. In this sense, Halit, de Haan, and Johnson (2000) report higher N170 amplitudes for atypical than for typical faces. In terms of facial expressions of emotion, meta-analytic evidence suggests that N170 is increased by emotional faces (Ferreira-Santos, Martins, Almeida, Barbosa, Marques-Teixeira, & de Haan, 2013; Hinojosa, Mercado, & Carretié, 2015). Additional results from our group suggest that this effect is driven by an arousal modulation, as we found N170 changes associated with the perceived arousal of facial expressions of emotion (FEE) regardless of emotional category or valence properties (Almeida, Ferreira-Santos, Chaves, Paiva, Barbosa & Marques-Teixeira, 2016; Ferreira-Santos, 2013). Additionally, the Vertex Positive Potential (VPP) is also considered one of the main ERP components in studying facial processing. The VPP consists of a positive deflection in fronto-central sites with a similar latency to N170, as well as a similar modulation by configural properties of the facial stimuli (Eimer, 2000) and presence of emotional content (Ashley, Vuillemier, & Swick, 2004). In fact, it is likely that both scalp potentials share the same neural dipole source and reflect the same processes (Joyce & Rossion, 2005; Luo, Feng, He, Wang, & Luo, 2010).

Considering the neuroimaging and electrophysiological evidence of both typicality and arousal modulations, we hypothesized that atypicality could have a potentially cumulative effect with arousal in the N170 and VPP. We also collected self-reported ratings of the facial stimuli both as manipulation checks and to examine if the associations in subjective reports mirror the electrophysiological results. Specifically, in terms of the subjective perception of the faces, we predicted that atypical stimuli would be also rated with more negative valence and higher arousal.

## Method

### Participants

Thirty adult participants (21 female, mean age = 24.0, *SD* = 3.87) were recruited from the community via website, mailing lists, and social media calls for study participation. All of them had normal or corrected to normal vision. Participants were screened for self-reported neurological and psychiatric disorders, head injuries, and substance abuse, which constituted exclusion criteria, but none were excluded for these reasons.

The local Ethics Committee approved the study and all participants provided written informed consent in accordance with the Declaration of Helsinki at the beginning of the experiment.

### Stimuli

A set of 80 pictures from the NimStim Face Stimulus Set (Tottenham et al., 2009) were selected based on previously obtained self-report ratings of typicality, valence, and arousal (see procedure below). Our goal was to select facial stimuli that differed in their affective properties within emotional categories (e.g., pleasant and unpleasant surprised faces, low and high arousal angry faces) for typical and atypical actors.

Pre-study typicality ratings were collected in a self-report pilot task with nine participants (five female; mean age = 25.5, *SD* = 4.56). Participants were required to rate the typicality of each actor’s neutral expression in a 7-point Likert scale. Based on these data, we selected the most typical and atypical actors from the NimStim dataset. The choice of the neutral expression to obtain typicality ratings followed from the fact that it is likely the best image to evaluate face typicality in terms of identity and facial structure, without being confounded by the typicality/frequency of the facial expression (e.g., happy facial expressions have been reported to be more typical/frequent than others; Sommerville & Whalen, 2006). The average value of (prestudy) typicality was calculated for each actor, and the most (> Q3) and the least (< Q1) typical actors were selected. This led to a total of eight atypical actors (three female) and nine typical actors (five female).

As specific emotions are collinear with levels of valence or arousal (e.g., anger expressions are necessarily unpleasant and neutral/calm faces are always low in arousal), dissociation of arousal and valence within emotional categories can only be achieved in a partial factorial design (Almeida et al., 2016; Ferreira-Santos, 2013). Here we contrasted Low and High Arousal exemplars of Angry and Happy expressions and Pleasant and Unpleasant exemplars of Calm and Surprised expressions. To select these stimuli, we considered the actors selected as the most typical or atypical, and extracted the ratings of valence for their Calm and Surprised expressions, and ratings of arousal for their Angry and Happy expressions from a reference database (*N* = 40, young adults, 20 female) obtained in a previous study (Ferreira-Santos, 2013). We then calculated the 35^th^ and 65^th^ percentiles for valence and for arousal, and selected the images that were more extreme than those values: the most pleasant and unpleasant images of Calm and Surprise, and the lowest and highest arousal images of Anger and Happiness. Table 1 summarizes the experimental conditions and selection criteria (and the codes of the actual NimStim stimuli used are indicated in Footnote 1^1^).

**Table 1.** Experimental conditions resulting from NimStim stimuli selection criteria based on levels of emotional category, valence/arousal, and typicality.

| Emotion | Affect | Typicality |  |
| --- | --- | --- | --- |
|  |  | Typicality ratings > Q3 | Typicality ratings < Q1 |
| Calm | Valence < 35<sup>th</sup> percentile | Typical Calm Unpleasant | Atypical Calm Unpleasant |
|  | Valence > 65<sup>th</sup> percentile | Typical Calm Pleasant | Atypical Calm Pleasant |
| Surprise | Valence < 35<sup>th</sup> percentile | Typical Surprise Unpleasant | Atypical Surprise Unpleasant |
|  | Valence > 65<sup>th</sup> percentile | Typical Surprise Pleasant | Atypical Surprise Pleasant |
| Anger | Arousal < 35<sup>th</sup> percentile | Typical Anger Low Arousal | Atypical Anger Low Arousal |
|  | Arousal > 65<sup>th</sup> percentile | Typical Anger High Arousal | Atypical Anger High Arousal |
| Happiness | Arousal < 35<sup>th</sup> percentile | Typical Happiness Low Arousal | Atypical Happiness Low Arousal |
|  | Arousal > 65<sup>th</sup> percentile | Typical Happiness High Arousal | Atypical Happiness High Arousal |

<sup>1</sup> Atypical Happiness Low Arousal: 18F\_HA\_C, 19F\_HA\_C, 26M\_HA\_C, 38M\_HA\_C, 45M\_HA\_C; Atypical Happiness High Arousal: 13F\_HA\_X, 19F\_HA\_X, 22M\_HA\_X, 37M\_HA\_X, 45M\_HA\_X; Typical Happiness Low Arousal: 03F\_HA\_C, 09F\_HA\_C, 20M\_HA\_C, 24M\_HA\_C, 29M\_HA\_C; Typical Happiness High Arousal: 08F\_HA\_X, 09F\_HA\_X, 27M\_HA\_X, 29M\_HA\_X, 36M\_HA\_X; Atypical Anger Low Arousal: 15F\_AN\_C, 18F\_AN\_C, 19F\_AN\_C, 31M\_AN\_C, 32M\_AN\_C; Atypical Anger High Arousal: 14F\_AN\_O, 31M\_AN\_O, 37M\_AN\_O, 38M\_AN\_O, 45M\_AN\_O; Typical Anger Low Arousal: 03F\_AN\_C, 06F\_AN\_C, 24M\_AN\_C, 28M\_AN\_C, 29M\_AN\_C; Typical Anger High Arousal: 03F\_AN\_O, 10F\_AN\_O, 29M\_AN\_O, 34M\_AN\_O, 36M\_AN\_O; Atypical Calm Unpleasant: 02F\_CA\_O, 13F\_CA\_O, 14F\_CA\_O, 19F\_CA\_C, 32M\_CA\_O; Atypical Calm Pleasant: 15F\_CA\_C, 18F\_CA\_C, 22M\_CA\_C, 32M\_CA\_C, 45M\_CA\_C; Typical Calm Unpleasant: 06F\_CA\_O, 09F\_CA\_O, 20M\_CA\_O, 29M\_CA\_O, 43M\_CA\_O; Typical Calm Pleasant: 03F\_CA\_O, 11F\_CA\_C, 24M\_CA\_C, 28M\_CA\_C, 39M\_CA\_C; Atypical Surprise Unpleasant: 13F\_SP\_O, 15F\_SP\_O, 30M\_SP\_O, 25M\_SP\_O, 38M\_SP\_O; Atypical Surprise Pleasant: 02F\_SP\_O, 12F\_SP\_O, 15F\_SP\_C, 22M\_SP\_O, 37M\_SP\_O; Typical Surprise Unpleasant: 06F\_SP\_O, 08F\_SP\_O, 09F\_SP\_O, 39M\_SP\_O, 43M\_SP\_O; Typical Surprise Pleasant: 03F\_SP\_O, 10F\_SP\_O, 20M\_SP\_O, 27M\_SP\_O, 34M\_SP\_O.

Each condition, as displayed in Figure 1, included five stimuli. The valence and arousal values for each condition did not significantly differ between atypical and typical groups (paired t-tests).

**Figure 1.**
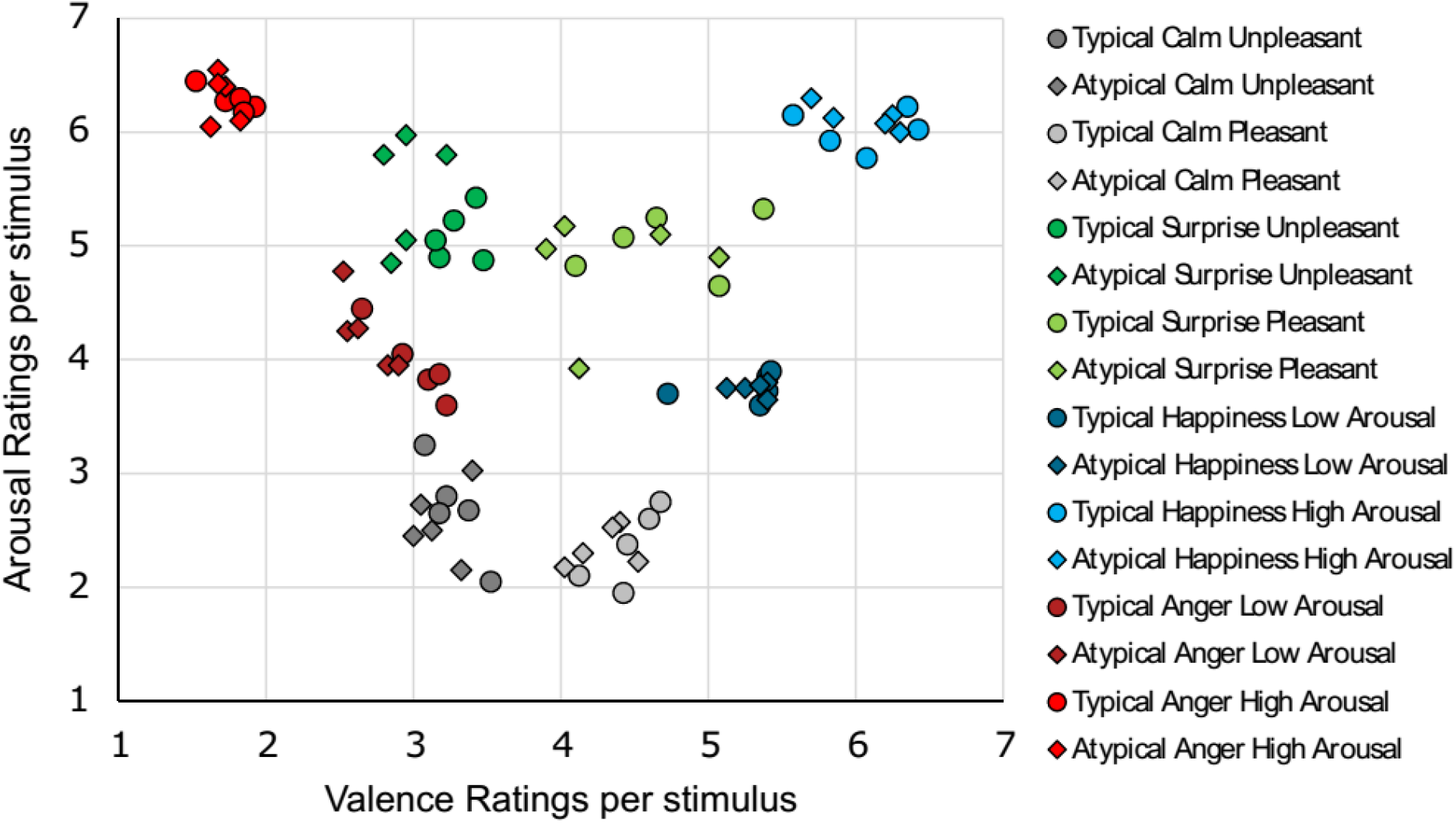
Scatterplot of valence and arousal ratings for the selected stimuli (from reference data; Ferreira-Santos, 2013).

### Procedure

After signing the consent form and preparation of the EEG apparatus (described below), participants were comfortably seated in a data collection chamber. The study was composed by an EEG task followed by a ratings task, in this order for all participants. In the EEG task, participants were asked to press a button on a controller whenever they saw an inverted face. These target inverted faces were only presented to ensure participants remained attentive throughout the experiment. Each trial began with a fixation cross in the centre of the screen for a variable duration (1000-1500 ms), followed by a facial stimulus presented for 500 ms. The experiment had a total of 440 trials: 16 conditions x 25 trials (as each stimulus was presented 5 times), plus 40 target inverted faces. The trials were equally divided in two blocks with a short break between them, to avoid fatigue. After the EEG task was completed, participants were asked to rate each of the stimuli used in the EEG task for arousal, valence, and typicality in a 7-point Likert scale and to identify the emotional category represented from a list of basic emotions/emotional categories (anger, surprise, fear, sadness, happiness, disgust, neutral, calm, and an option of “none of the others”). Both tasks were presented using E-Prime v2.0 (2011, Psychology Tools, Inc., Pittsburgh, PA, USA) in a 17’’ screen located 1.15 m from the participant, with facial stimuli subtending a horizontal and vertical visual angle of 7.46° by 11.91°, respectively. Stimuli in both tasks were presented in random order.

### EEG recording and signal pre-processing

EEG data were acquired at a digitizing rate of 500 Hz using a 128-channel EEG NetAmps system running NetStation v4.5.2 (2008, EGI, Electrical Geodesics, Inc., Eugene, OR, USA). All impedances were kept below 50 kOhm as this is a high impedance input amplifier. The signal was referenced online to the vertex (Cz).

EEG data were analysed using EEGLAB v13.6.5b (Delorme & Makeig, 2004), a MATLAB toolbox (2017, The Mathworks, Inc., Massachusetts, USA), and the ERPLAB plugin v6.1.3 (Lopez-Calderon & Luck, 2014). For ERP analysis, the data were band-pass filtered (0.1-30Hz). Some channels were permanently removed from analysis due to hardware problems (physically damaged electrodes in some of the sensor nets): midline electrodes were removed from the analysis if they were missing for more than half of the participants; for lateral channels, we additionally removed the symmetrical electrode when it was absent in more than 1/3 of the sample. According to these criteria, we excluded the following electrodes: E1, E6, E8, E17, E25, E32, E48, E119, E126, and E127. The resulting datasets contained a symmetrical montage of 119 channels. From these remaining electrodes, channels with bad data (flatline, excessive noise, or excessive drift) were removed. The signal was submitted to an Independent Component Analysis (ICA) to identify and correct eye blink and heart rate artefacts, followed by the removed channels’ interpolation. The signal was re-referenced to the average of all electrodes. Finally, epochs of 1000ms (-200ms pre-stimulus baseline) were visually inspected and those containing artefacts were rejected prior to averaging.

There were no significant differences between conditions in the final number of valid trials (repeated measures ANOVA with condition as factor (16 levels), *F*(15,435) = 1.24, *p* = .238, η^2^_p_ = .041). Based on the previous literature (Joyce & Rossion, 2005) and after a visual inspection of individual averages and grand-averages, the components of interest were extracted. Table 2 summarizes the measures that were extracted.

**Table 2.**
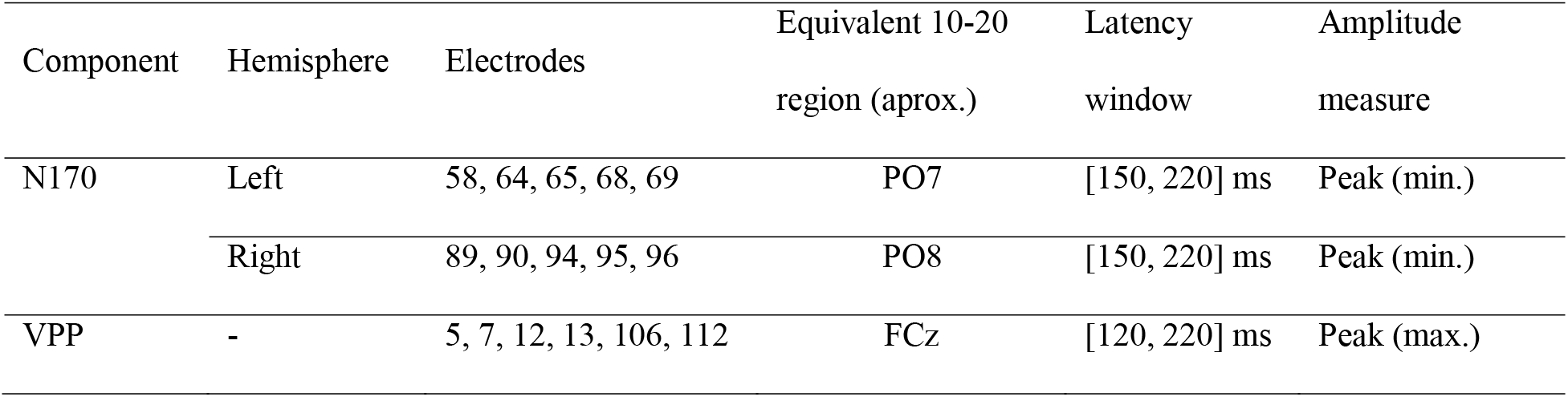
Summary of the parameters used to quantify the amplitudes of ERP components.

| Component | Hemisphere | Electrodes | Equivalent 10-20<br>region (aprox.) | Latency<br>window | Amplitude<br>measure |
| --- | --- | --- | --- | --- | --- |
| N170 | Left | 58, 64, 65, 68, 69 | PO7 | [150, 220] ms | Peak (min.) |
|  | Right | 89, 90, 94, 95, 96 | PO8 | [150, 220] ms | Peak (min.) |
| VPP | - | 5, 7, 12, 13, 106, 112 | FCz | [120, 220] ms | Peak (max.) |

### Statistical analysis

As a manipulation check to confirm that overall affective properties of the stimuli were perceived as expected, we compared the ratings of arousal and valence between the reference sample (used to select the stimuli) and the sample from the current study by means of t-tests.

In order to examine the effect of typicality, emotion, and the affective properties on our dependent variables for subjective ratings (ratings of arousal, valence, and typicality) and ERP measures (N170 and VPP amplitudes), separate Repeated Measures ANOVA were conducted in SPSS v24 (2016, IBM Statistics, New York, USA) for each outcome. Also, the factor for the affective properties varied: when comparing happy and angry expressions, the within-category affective property that was manipulated was the level of arousal; but when comparing calm and surprised faces, the affective property that varied within-categories was valence. As such, one set of ANOVAs had the following factors: *Typicality* (atypical, typical), *Emotion* (angry, happy), and *Arousal Level* (low arousal, high arousal). The other set of ANOVAs had *Typicality* (atypical, typical), *Emotion* (calm, surprise), and *Valence* (unpleasant, pleasant). The *Hemisphere* (left, right) was an additional factor for analysis of ERP measures recorded bilaterally.

The association between self-reported ratings and electrophysiological data including all conditions was estimated using multilevel models (MLM) to control for the dependencies in the data (as each participant provided multiple data points) in R 3.4.2 (R Core Team, 2017), using the nlme package (version 3.1-131; Pinheiro et al., 2017). We estimated separate models for N170 amplitude (averaged for both hemispheres) as a function of arousal, valence, or typicality ratings. Ratings were entered as a fixed effect and participants as random effects (random intercepts and slopes). Standardized coefficients for the association between N170 amplitude and ratings were calculated based on the fixed effect t-score estimates of the MLM.

The threshold for statistical significance for all analysis was set at α□ = □.05. Violations of sphericity were corrected by the Greenhouse-Geisser method and significant ANOVA main effects were quantified using Bonferroni-corrected post-hoc comparisons.

## Results

### Subjective ratings results

There were no significant differences concerning the ratings of arousal, valence, and typicality between the reference sample (used initially to select the stimuli) and the ratings provided by the sample of the present study (all *p*s > .05). Note that typicality ratings were not compared because in the pre-study these were only collected for neutral faces, most of which were not used in the current study. Figure 2 shows arousal and valence ratings given by the participants for each stimulus.

**Figure 2.**
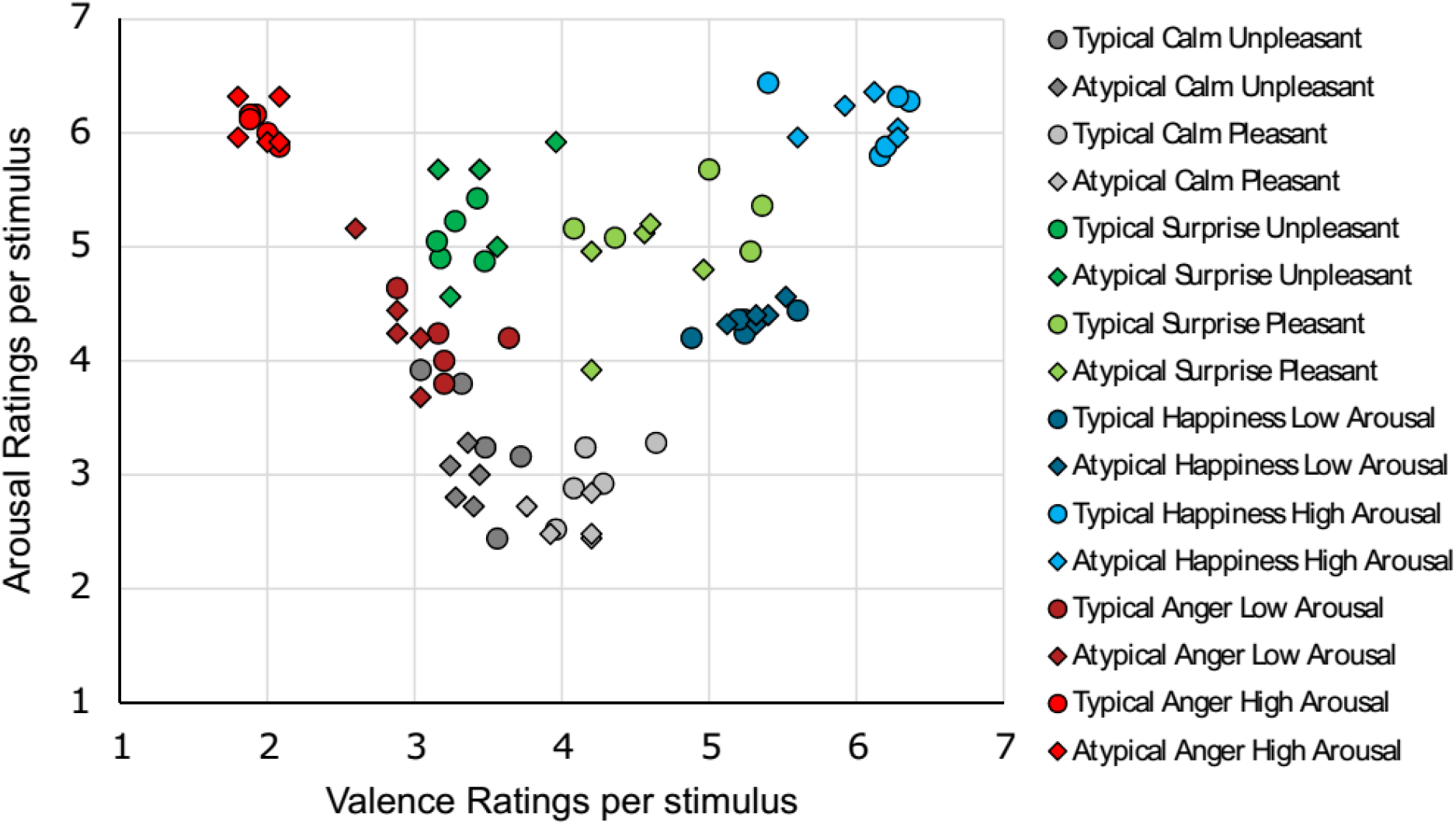
Scatterplot of valence and arousal ratings for the stimuli given by the participants in the present study.

### Arousal ratings

For the ANOVA involving Angry and Happy expressions, there was a main effect of *Arousal Level, F*(1,24) = 68.6, *p* < .001, η_p_^2^ = .741, where high arousal images elicit higher arousal ratings (*M* = 6.10, *SE* = 0.22) comparably to low arousal images (*M* = 4.31, *SE* = 0.16). There was no effect for *Emotion* or *Typicality* (*p* > .25). No significant first nor second order interactions were found.

For the ANOVA involving Calm and Surprised expressions, there was a main effect of *Typicality, F*(1,24) = 42.5, *p* < .001, η_p_^2^ = .639, where typical faces were rated with more arousal (*M* = 4.16, *SE* = 0.15) than the atypical (*M* = 3.93, *SE* = 0.15); a main effect of *Emotion, F*(1,24) = 97.2, *p* < .001, η_p_^2^ = .802, with higher arousal ratings for surprise (*M* = 5.14, *SE* = 0.16) than for calm (M= 2.96, *SE* = 0.21); and a main effect of *Valence, F*(1,24) = 7.12, *p* = .013, η_p_^2^ = .229, where unpleasant faces were rated as more aroused (*M* = 4.20, *SE* = 0.16) than pleasant (*M* = 3.90, *SE* = 0.15).

In terms of interactions, there was a significant *Typicality* by *Valence* interaction, *F*(1,24) = 13.6, *p* = .001, η_p_^2^ = .362, where typical pleasant expressions were rated as more aroused than the atypical (*p* < .001). Finally, there was a *Typicality* by *Emotion* by *Valence* interaction, *F*(1,24) = 8.80, *p* = .007, η_p_^2^ = .268.

### Valence ratings

The Angry-Happy ANOVA revealed a main effect of *Emotion, F*(1,24) = 119, *p* < .001, η_p_^2^ = .832, with more positive ratings of valence to expressions of happiness (*M* = 5.67, *SE* = 0.11) comparably to faces of anger (*M*= 2.50, *SE* = 0.21); and a main effect of *Arousal Level, F*(1,24) = 6.40, *p* = .018, η_p_^2^ = .210, where low arousal images (*M* = 4.17, *SE* = 0.10) elicited more positive ratings of valence (*M*= 4.01, *SE* = 0.08).

There was a significant *Emotion* by *Arousal Level* interaction, *F*(1,24) = 46.4, *p* < .001, η_p_^2^ = .659, with differences between all the conditions (*p* < .001).

Finally, there was a *Typicality* by *Emotion* by *Arousal Level* interaction, *F*(1,24) = 10.3, *p* = .004, η_p_^2^ = .301. Here, for low arousal anger expressions, there were more positive ratings of valence for the typical than for the atypical faces (*p* < .001).

For the Calm-Surprise ANOVA we found a main effect of *Typicality, F*(1,24) = 17.7, *p* < .001, η_p_^2^ = .424, with more positive ratings for typical (*M* = 4.03, *SE* = 0.10) than for atypical actors (*M* = 3.84, *SE* = 0.10); a main effect of *Emotion, F*(1,24) = 6.56, *p* = .016, η_p_^2^ = .217, with more positive ratings for surprise (*M* = 4.11, *SE* = 0.13) than for calm (*M* = 3.77, *SE* = 0.96); and a main effect of *Valence, F*(1,24) = 37.1, *p* < .001, η_p_^2^ = .607, with higher valence ratings for pleasant (*M* = 4.40, *SE* = 0.09) than for unpleasant faces (*M*= 3.47, *SE* = 0.15).

The only significant interaction was *Emotion* by *Valence, F*(1,24) = 6.81, *p* = .015, η_p_^2^ = .221, where, for pleasant expressions, there were more positive ratings for surprised than for calm faces (*p* < .001).

### Typicality ratings

The Angry-Happy ANOVA revealed a main effect of *Typicality, F*(1,24) = 4.82, *p* = .038, η_p_^2^ = .167, with higher ratings of typicality for the typical faces (*M* = 4.83, *SE* = 0.17) comparably to the atypical (*M* = 4.56, *SE* = 0.20), as well as an *Emotion* main effect, *F*(1,24) = 39.2, *p* < .001, η_p_^2^ = .620, where happy faces were rated as more typical (*M*= 5.30, *SE* = 0.18) than angry expressions (*M* = 4.09, *SE* = 0.22), and an *Arousal Level* main effect, *F*(1,24) = 4.63, *p* = .042, η_p_^2^ = .162, with low arousal expressions eliciting higher ratings of typicality (*M* = 4.91, *SE* = 0.20) than high arousal faces (*M* = 4.47, *SE* = 0.21).

Also, there was a significant *Typicality* by *Emotion* by *Arousal Level* interaction, *F*(1,24) = 4.74, *p* = .040, η_p_^2^ = .165. No other significant interactions were found.

Concerning the Calm-Surprise ANOVA, we obtained a main effect of *Typicality, F*(1,24) = 7.36, *p* = .012, η_p_^2^ = .235, with higher typicality ratings for typical faces (*M* = 4.61, *SE* = 0.15) comparably to atypical (*M*= 4.38, *SE* = 0.17), and a *Valence* main effect *F*(1,24) = 31.0, *p* < .001, η_p_^2^ = .564, where pleasant expressions were rated as more typical (*M*= 4.78, *SE* = 0.16) comparably to unpleasant expressions (*M* = 4.21, *SE* = 0.16).

A significant interaction between *Emotion* and *Valence* was also found, *F*(1,24) = 5.98, *p* = .022, η_p_^2^ = .199, where for both calm (typical, *M*=4.89, *SE* = 1.01, and atypical, *M* = 4.82, *SE* = 1.21, *p* < .001) and surprise (typical, *M* = 4.79, *SE* = 0.99, and atypical, *M* = 4.61, *SE* = 0.97, *p* = .002) the pleasant images were rated as more typical than for unpleasant configurations of calm (typical, *M* = 4.17, *SE* = 0.90, and atypical, *M* = 3.92, *SE* = 0.99) and surprise (typical *M* = 4.58, *SE* = 1.14, and atypical, *M* = 4.18, *SE* = 1.11). No other interactions were significant.

### ERP results N170

#### Angry-Happy ANOVA

There was a main effect of *Arousal Level, F*(1,29) = 11.4, *p* = .002, η_p_^2^= .282, where high arousal expressions showed larger N170 amplitudes (*M* = -2.36, *SE* = 0.37) than low arousal images (*M* = -1.87, *SE* = 0.36). Also, there was a significant *Hemisphere* by *Emotion* interaction, *F*(1,29) = 5.28, *p* = .029, η_p_^2^= .154, where, for anger, there were larger amplitudes in the right (*M* = -2.52, *SE* = 0.46) than on the left hemisphere (*M* = −1.71, *SE* = 0.35, *p* = .045) (Figure 3).

**Figure 3.**
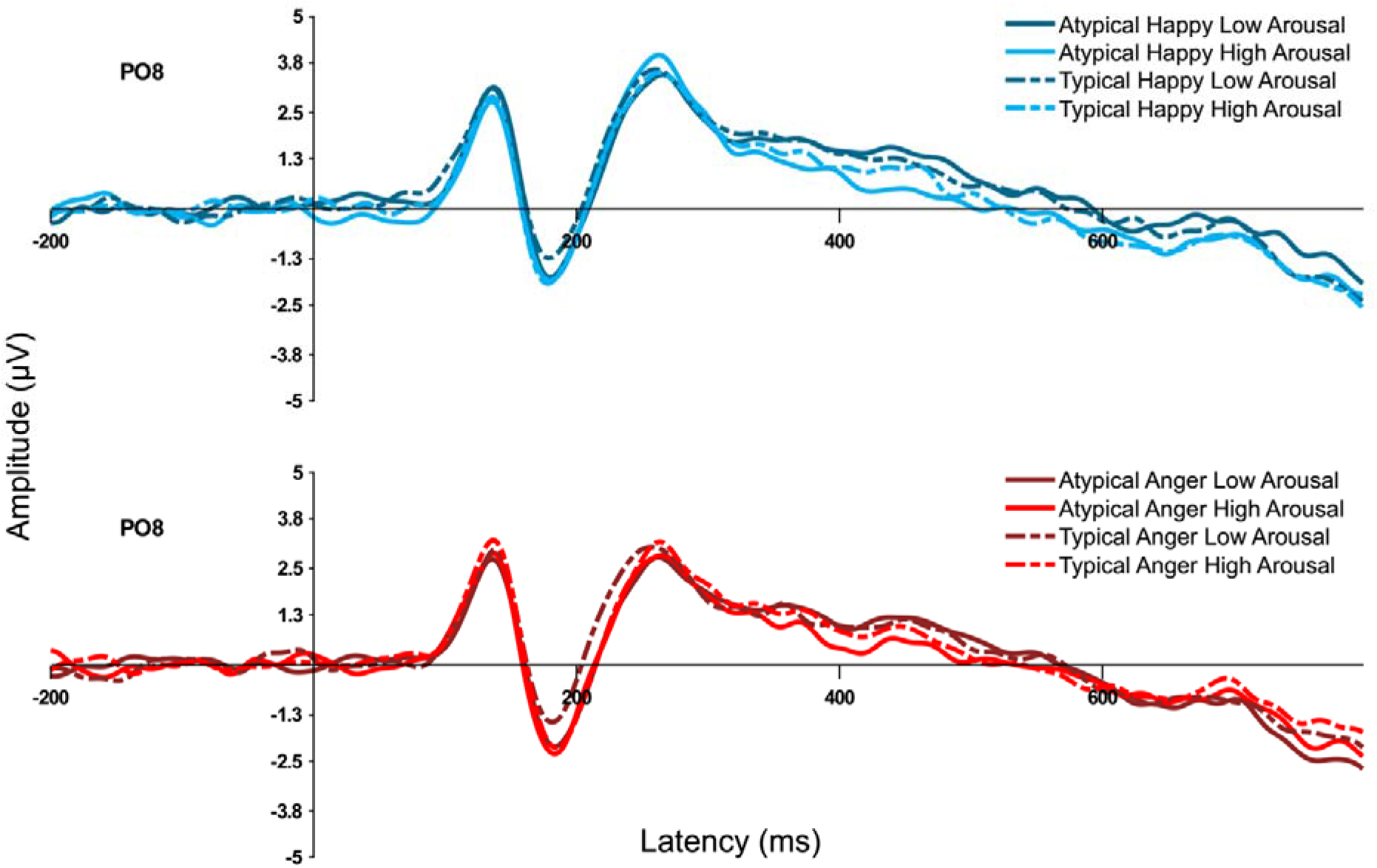
Illustrative ERP waveform at PO8 for the Angry/Happy analysis.

### Calm-Surprise ANOVA

There was a main effect of *Hemisphere, F*(1,29) = 5.34, *p* = .028, η_p_^2^= .155, with larger amplitudes in right sites (*M* = -2.45, *SE* = 0.43) than for the left side (*M* = -1.67, *SE* = 0.34); a main effect of *Typicality, F*(1,29) = 11.1, *p* = .002, η_p_^2^ = .276, with higher amplitudes to typical faces (*M* = -2.23, *SE* = 0.36) comparably to the atypical (*M* = -1.88, *SE* = 0.35); a main effect of *Emotion, F*(1,29) = 4.91, *p* = .035, η_p_^2^= .145, with larger amplitudes for surprise (*M* = -2.20, *SE* = 0.34) than for expressions of calm (*M*= -1.92, *SE* =0.37); and also a *Valence* main effect, *F*(1,29) = 12.4, *p* = .001, ηp^2^ = .299, with higher amplitudes for unpleasant facial expressions (*M* = -2.25, *SE* = 0.36) than pleasant (*M* = -1.86, *SE* = 0.35) (Figure 4).

**Figure 4.**
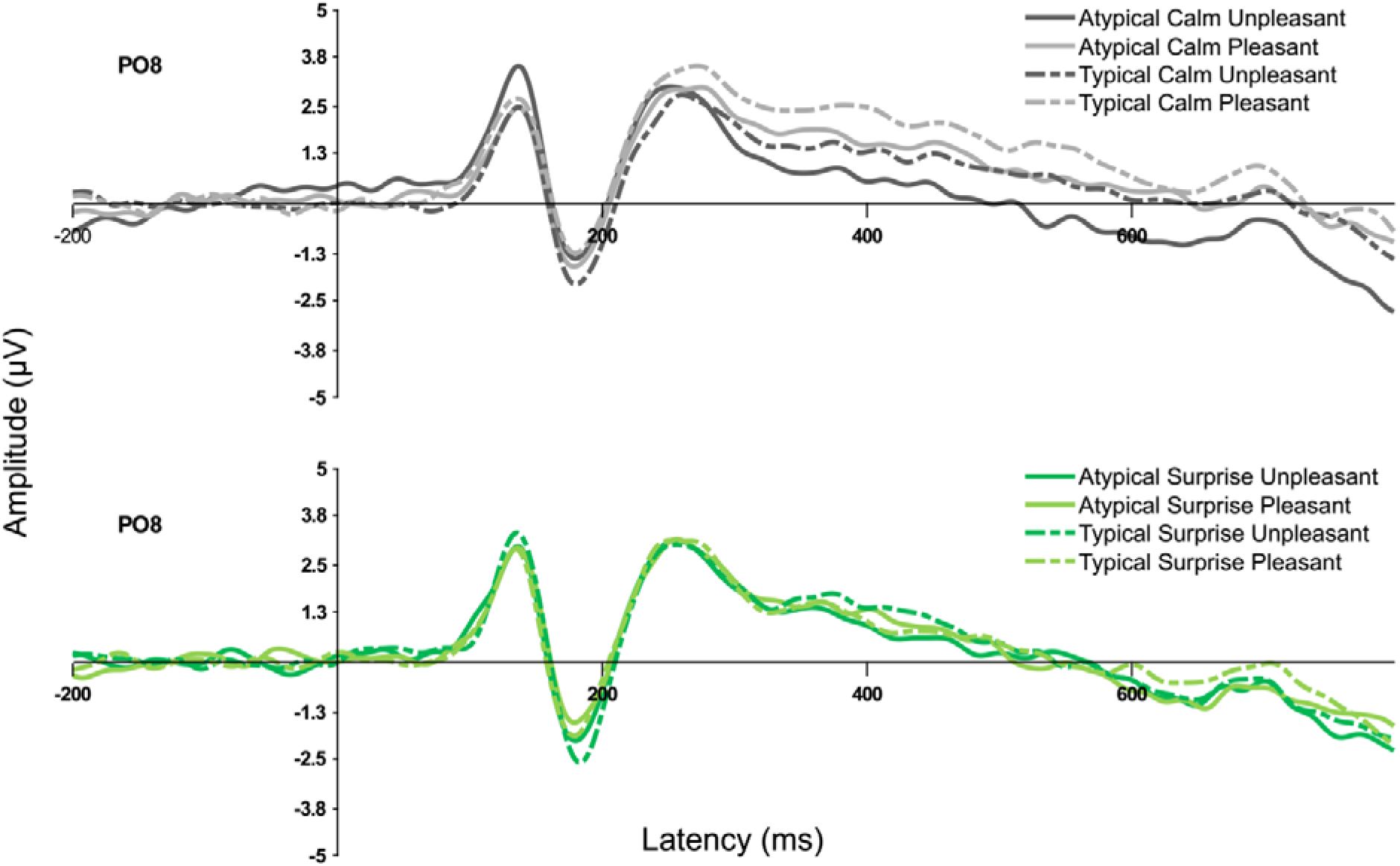
Illustrative ERP waveform at PO8 for the Calm/Surprise analysis.

No significant interactions were found.

### VPP

#### Angry-Happy ANOVA

There was only one significant effect, namely of *Arousal Level, F*(1,29) = 14.9, *p* = .001, η_p_^2^ = .339, with larger amplitudes for high arousal facial expressions (*M* = 2.02, *SE* = 0.29) than for low arousal expressions (*M* = 1.49, *SE* = 0.28).

#### Calm-Surprise ANOVA

There was a significant *Typicality* by *Valence* interaction, *F*(1,29) = 5.87, *p* = .022, η_p_^2^ = .168, where for unpleasant pictures, typical actors elicited higher amplitudes (*M* = 2.27, *SE* = 0.32) than atypical (*M* = 1.77, *SE* = 0.29) (*p* = .016).

### Association between ratings and ERP results

Analysis via multilevel models revealed a modest but significant association between the N170 amplitude and arousal ratings, β = .111, *p* = .031, where larger N170 amplitudes (i.e., more negative amplitudes) were associated with higher arousal ratings. Valence and typicality were not significantly associated with ERP responses.

## Discussion

In the present study, we aimed to dissociate the impact of face typicality and affective properties in emotional face processing. To this end, we presented typical and atypical actors performing facial expressions of emotion that varied in arousal and valence. We collected subjective ratings of valence, arousal, and typicality as well as neural responses to these stimuli, specifically the N170 and VPP.

First, the subjective ratings provided a manipulation check for our design, confirming that the experimental conditions were generally in accordance to our stimuli selection. Additionally, individual ratings shed light on the interaction between affective properties and face typicality. We will first discuss the results for ratings of arousal, valence, and typicality, and do so separately for each analysis (the Angry/Happy analysis, where arousal level within each category was manipulated and the Calm/Surprise analysis, where valence was manipulated).

In the Angry/Happy analysis, high arousal images elicited (unsurprisingly) higher arousal ratings, but no main effects of nor interaction with emotion or typicality were found. This finding does not support our hypothesis that atypical faces would elicit higher ratings of arousal.

In the Calm/Surprise analysis, the pattern of results was more complex as we found higher arousal ratings for Surprise expressions, for actors with typical and unpleasant faces. Firstly, the attribution of higher arousal ratings for Surprise is in line with the previous arousal hypothesis, since this emotional category is characterized by high levels of activation in both Russell’s circumplex model of emotion (Russell, 1980) and our reference data (Ferreira-Santos, 2013). The typicality effect was unexpected, but it may reflect the findings of Sánchez and Vasquéz (2013), where the prototypicality of the emotional face was associated with higher perceived activation. This is also corroborated by the higher arousal ratings attributed to typical than atypical pleasant expressions. Finally, the attribution of more perceived arousal to unpleasant expressions is not new in the literature (Adolphs, Russell, & Tranel, 1999).

In sum, regarding the arousal ratings, images previously selected based on higher prestudy arousal levels (meaning high arousal stimuli for Angry/Happy and surprise stimuli for Calm/Surprise) and unpleasant stimuli were subjectively rated as more aroused, as expected, and typicality and emotional category did not seem to affect the subjective ratings of arousal.

In terms of perceived valence, in the Angry/Happy analysis, more positive valence was attributed to happy faces, as expected. Also, low arousal images were perceived as more pleasant. A further interaction between *Emotion* and *Arousal Level* contributed to this effect: low-arousal anger expressions and high-arousal happy expressions were the most positively rated. In the Calm/Surprise analysis, pleasant faces of both emotional categories received more positive ratings, as expected. Surprised faces were also considered more pleasant than calm expressions. However, the dispersion of valence ratings was similar for both emotional categories, with a few surprised faces being rated as very pleasant. Typical faces are again rated as more pleasant, which is consistent with the literature mentioned above about positive affect (Lee, 2001).

Finally, typicality ratings were higher for typical faces in both analyses, confirming our design. In the Angry/Happy analysis, happy expressions and low arousal stimuli were considered more typical. In line with these findings is the result found in the Calm/Surprise analysis, in which the more pleasant the face, the more typical it was considered.

These results allowed to confirm that our experimental manipulation was successful in both analyses (Angry/Happy and Calm/Surprise) but also to draw conclusions about the relation between typicality and the affective properties of arousal and valence. Specifically, low arousal and more pleasant faces seem to be considered more typical. If we consider the face-space model and the concept of “prototypical face” as the average of all the sampled faces (Valentine, 1991; Valentine & Ferrara, 1991), we can assume that the low-arousal form is indeed the most prototypical for each emotional expression. Also, the association between typicality, pleasantness, and happiness could be due to the fact that positive expressions are the most common expression in the life-span of healthy young adults, comparably to negative expressions such as anger and fear (Somerville & Whalen, 2006). This effect is also well described in low-risk developmental samples as the “happiness advantage” (de Haan & Matheson, 2009). From a Predictive Processing perspective, this would perhaps suggest that the overall prior for emotional expressions consists of low arousal pleasant expressions.

Looking at the ERP results, N170 and VPP amplitudes were modulated by arousal in both analyses. This arousal modulation was expected based on previous work from our group (Almeida et al., 2016; Ferreira-Santos, 2013), but the hypothesized modulation by typicality was not found. The effects of emotion (Calm, Surprise) on the N170 were more complex.

Expressions of surprise evoked higher amplitudes than calm faces, which can be considered an effect of arousal. Also, unpleasant faces induced higher N170 than pleasant, as did typical faces comparably to atypical, which is corroborated by the VPP results with higher amplitudes for typical unpleasant. However, this pattern of significant differences becomes clear when compared with the self-reported arousal ratings for Calm and Surprised faces. Indeed, the same pattern of effects was found for arousal ratings and the N170 amplitude, again supporting arousal as a crucial affective property influencing the electrophysiological response to facial expressions of emotions. Indeed, an association between arousal ratings and N170 amplitude was the only statistically significant effect found in the multilevel model analysis. The magnitude of this effect was modest, but it should be noted that this effect reflects a modulation of ERP amplitude among several sources of variance. It is known that N170 amplitude is primarily influenced by the presence of a face stimulus and additional visual aspects additionally modulate the component amplitude, but in a much smaller degree (Rossion et al., 2000). In the present study stimuli varied not only in arousal but also in actor identity and gender, emotional category, valence, and typicality, but only arousal significantly predicted N170 amplitude. Thus, aspects as typicality or valence may only modulate electrophysiological patterns if they also affect the perceived arousal of the emotional faces.

Another outstanding issue, and a limitation of the present work, is how to define the affective concepts we are working with beyond their use in self-reports. Valence is perhaps more straight-forward as it can be linked to experiences and representations of rewards and punishments (Bach & Dayan, 2017) but a definition of arousal is less clear. In facial expressions, arousal relates to the intensity of the facial display and thus co-varies with perceptual features of the stimuli that signal facial muscle activity (e.g., open mouth, skin wrinkles, relative proportion of visible sclera in the eyes; Calvo & Nummenmaa, 2016; Whalen et al., 2004). However, affective arousal is likely a more complex construct that involves multiple autonomic and central nervous system components and may play an important role not only in intense emotional responses (where it has been best documented) but also in general cortical function by modulating the processing and memorization of stimuli (Ferreira-Santos, 2016; Mather, Clewett, Sakaki, & Harley, 2016; Satpute, Kragel, Barrett, Wager, & Bianciardi, 2018).

Additionally, these results help us rethink the concept of typicality in emotional face processing. If face typicality is considered the proximity to the average configuration postulated by the Face-Space model, it may also be considered the similarity to the prior of a face, in a predictive processing perspective. In this sense, in cases of neutral facial expressions, as used in the studies presented above regarding trustworthiness and attractiveness, the prediction error will be evoked and proportional to the distance between the facial configuration and the neutral prior, thus eliciting higher neural processing the more atypical the face stimulus is. In the case of emotional face processing, further research is needed in the field of Predictive Processing to understand how an emotional face prior may be developed. Clark (2013) considers that a prior is built in a three-stage process, evolving from an abundance of prediction errors, to a convergence to a general theme, that is later on negotiated and sensible to perceptual and contextual changes at each contact with the world. In this sense, at each contact with emotional faces, the existence of “subface spaces” for each emotional category is possible, each with their own prototype (i.e., average configuration). In this sense, contrarily to the processing of neutral faces, changes in the activation or arousal of emotional faces may be the main modulators of brain activation by being the perceptual and affective evokers of prediction error. Thereby, arousal may change the level of typicality of an emotional face, and not the other way around. However, further research on emotional face processing under a predictive processing approach is needed.

The results of this work corroborate the findings on the central role of arousal in emotional face processing (Almeida et al., 2016) and may provide a challenge to the typicality explanation of the non-linear results found in neuroimaging studies. Further research that considers both affective and typicality factors in neuroimaging designs could provide additional evidence to the disentanglement of these two properties.

## Footnotes

1 Atypical Happiness Low Arousal: 18F_HA_C, 19F_HA_C, 26M_HA_C, 38M_HA_C, 45M_HA_C; Atypical Happiness High Arousal: 13F_HA_X, 19F_HA_X, 22M_HA_X, 37M_HA_X, 45M_HA_X; Typical Happiness Low Arousal: 03F_HA_C, 09F_HA_C, 20M_HA_C, 24M_HA_C, 29M_HA_C; Typical Happiness High Arousal: 08F_HA_X, 09F_HA_X, 27M_HA_X, 29M_HA_X, 36M_HA_X; Atypical Anger Low Arousal: 15F_AN_C, 18F_AN_C, 19F_AN_C, 31M_AN_C, 32M_AN_C; Atypical Anger High Arousal: 14F_AN_O, 31M_AN_O, 37M_AN_O, 38M_AN_O, 45M_AN_O; Typical Anger Low Arousal: 03F_AN_C, 06F_AN_C, 24M_AN_C, 28M_AN_C, 29M_AN_C; Typical Anger High Arousal: 03F_AN_O, 10F_AN_O, 29M_AN_O, 34M_AN_O, 36M_AN_O; Atypical Calm Unpleasant: 02F_CA_O, 13F_CA_O, 14F_CA_O, 19F_CA_C, 32M_CA_O; Atypical Calm Pleasant: 15F_CA_C, 18F_CA_C, 22M_CA_C, 32M_CA_C, 45M_CA_C; Typical Calm Unpleasant: 06F_CA_O, 09F_CA_O, 20M_CA_O, 29M_CA_O, 43M_CA_O; Typical Calm Pleasant: 03F_CA_O, 11F_CA_C, 24M_CA_C, 28M_CA_C, 39M_CA_C; Atypical Surprise Unpleasant: 13F_SP_O, 15F_SP_O, 30M_SP_O, 25M_SP_O, 38M_SP_O; Atypical Surprise Pleasant: 02F_SP_O, 12F_SP_O, 15F_SP_C, 22M_SP_O, 37M_SP_O; Typical Surprise Unpleasant: 06F_SP_O, 08F_SP_O, 09F_SP_O, 39M_SP_O, 43M_SP_O; Typical Surprise Pleasant: 03F_SP_O, 10F_SP_O, 20M_SP_O, 27M_SP_O, 34M_SP_O.

## References

Adolphs, R., Russell, J. A., & Tranel, D. (1999). A role for the human amygdala in recognizing emotional arousal from unpleasant stimuli. Psychological Science, 10, 167–171. https://doi.org/10.1111/1467-9280.00126

Almeida, P. R., Ferreira-Santos, F., Chaves, P. L., Paiva, T. O., Barbosa, F., & Marques-Teixeira, J. (2016). Perceived arousal of facial expressions of emotion modulates the N170, regardless of emotional category: Time domain and time-frequency dynamics. International Journal of Psychophysiology, 99, 48–56. https://doi.org/10.1016/j.ijpsycho.2015.11.017

Anderson, A. K., Christoff, K., Stappen, I., Panitz, D., Ghahremani, D. G., Glover, G., Gabrieli, J.D., & Sobel. N. (2003). Dissociated neural representations of intensity and valence in human olfaction. Nature Neuroscience, 6, 196–202. https://doi.org/10.1038/nn1001

Arnal, L. H., & Giraud, A.-L. (2012). Cortical oscillations and sensory predictions. Trends in Cognitive Sciences, 16, 390–398. https://doi.org/10.1016/j.tics.2012.05.003

Ashley, V., Vuilleumier, P., & Swick, D. (2004). Time course and specificity of event-related potentials to emotional expressions. NeuroReport, 15, 211–216.

Bach, D. R., & Dayan, P. (2017). Algorithms for survival: A comparative perspective on emotions. Nature Reviews Neuroscience, 18, 311–319. https://doi.org/10.1038/nrn.2017.35

Bentin, S., Allison, T., Puce, A., Perez, E., & McCarthy, G. (1996). Electrophysiological studies of face perception in humans. Journal of Cognitive Neuroscience, 8, 551–565. https://dx.doi.org/10.1162%2Fjocn.1996.8.6.551

Brodski-Guerniero, A., Paasch, G.-F., Wollstadt, P., Özdemir, I., Lizier, J. T., & Wibral, M. (2017). Information-theoretic evidence for Predictive Coding in the face-processing system. Journal of Neuroscience, 37, 8273–8283. https://doi.org/10.1523/JNEUROSCI.0614-17.2017

Bruce, V., & Young, A. (1986). Understanding race recognition. British Journal of Psychology, 77, 305–327. https://doi.org/10.1111/j.2044-8295.1986.tb02199.x

Calvo, M. G., & Nummenmaa, L. (2016). Perceptual and affective mechanisms in facial expression recognition: An integrative review. Cognition and Emotion, 30, 1081–1106. https://doi.org/10.1080/02699931.2015.1049124

Chatterjee, A., Thomas, A., Smith, S. E., & Aguirre, G. K. (2009). The neural response to facial attractiveness. Neuropsychology, 23, 135–143. https://doi.org/10.1037/a0014430

de Haan, M., & Matheson, A. (2009). The development and neural bases of processing emotion in faces and voices. In M. de Haan & M. R. Gunnar (Eds.), Handbook of Developmental Social Neuroscience (pp. 107–121). New York: Guilford Press.

DeBruine, L. M., Jones, B. C., Unger, L., Little, A. C., & Feinberg, D. R. (2007). Dissociating averageness and attractiveness: Attractive faces are not always average. Journal of Experimental Psychology: Human Perception and Performance, 33, 1420–1430. https://doi.org/10.1037/0096-1523.33.6.1420

Diamond, R., & Carey, S. (1986). Why faces are and are not special: An effect of expertise. Journal of Experimental Psychology: General, 115, 107–117. https://doi.org/10.1037/0096-3445.115.2.107

Eimer, M. (2000). The face-specific N170 component reflects late stages in the structural encoding of faces. Neuroreport, 11, 2319–2324.

Faerber, S.J., Kaufmann, J. M., Leder, H., Martin, E.-M., & Schweinberger, S. R. (2016). The role of familiarity for representations in norm-based face space. PLOS ONE, 11: e0155380. https://doi.org/10.1371/journal.pone.0155380

Ferreira-Santos, F. (2013). Modulation of event-related potentials by facial expressions of emotion in infants (at 9, 16, and 24 months) and adults: contributions for the understanding of the ontogenesis of emotional face processing (Doctoral thesis). University of Porto, Portugal.

Ferreira-Santos, F. (2016). The role of arousal in predictive coding [Commentary]. Behavioral and Brain Sciences, 39, e207. https://doi.org/10.1017/S0140525X15001788

Ferreira-Santos, F., Martins, E. C, Almeida, P. R., Barbosa, F., Marques-Teixeira, J., & de Haan, M. (2013). O processamento electrocortical de face é potenciado por expressões emocionais: Meta-análise do efeito de expressões faciais de emoção no componente N170 [Electrocortical face processing is enhanced by emotional expressions: Meta-analysis of the effect of facial expressions of emotion on the N170 component]. In A. Pereira, M. Calheiros, P. Vagos, I. Direito, S. Monteiro, C. F. Silva & A. A. Gomes (Eds.), Livro de Atas: VIII Simpósio Nacional de Investigação em Psicologia (pp. 857–866). Aveiro: Associação Portuguesa de Psicologia.

Hadj-Bouziane, F., Liu, N., Bell, A. H., Gothard, K. M., Luh, W.-M., Tootell, R. B. H.,… Ungerleider, L. G. (2012). Amygdala lesions disrupt modulation of functional MRI activity evoked by facial expression in the monkey inferior temporal cortex. Proceedings of the National Academy of Sciences of the United States of America, 109, E3640–E3648. https://doi.org/10.1073/pnas.1218406109

Halit, H., de Haan, M., & Johnson, M. H. (2000). Modulation of event-related potentials by prototypical and atypical faces. Neuroreport, 11, 1871–1875.

Hinojosa, J. A., Mercado, F., & Carretié, L. (2015). N170 sensitivity to facial expression: A meta-analysis. Neuroscience & Biobehavioral Reviews, 55, 498–509. https://doi.org/10.1016/j.neubiorev.2015.06.002

Iaria, G., Fox, C. J., Waite, C. T., Aharon, I., & Barton, J. J. S. (2008). The contribution of the fusiform gyrus and superior temporal sulcus in processing facial attractiveness: Neuropsychological and neuroimaging evidence. Neuroscience, 155, 409–422. https://doi.org/10.1016/j.neuroscience.2008.05.046

Joyce, C., & Rossion, B. (2005). The face-sensitive N170 and VPP components manifest the same brain processes: the effect of reference electrode site. Clinical Neurophysiology, 116, 2613–2631. https://doi.org/10.1016/j.clinph.2005.07.005

Kanwisher, N., & Yovel, G. (2006). The fusiform face area: A cortical region specialized for the perception of faces. Philosophical Transactions of the Royal Society B: Biological Sciences, 361(1476), 2109–2128. http://doi.org/10.1098/rstb.2006.1934

Kensinger, E. A., & Schacter, D. L. (2006). Processing emotional pictures and words: Effects of valence and arousal. Cognitive, Affective and Behavioural Neuroscience, 6, 110–126. https://doi.org/10.3758/CABN.6.2.110

Langlois, J. H., & Roggman, L. A. (1990). Attractive faces are only average. Psychological Science, 1, 115–122. https://doi.org/10.1111/j.1467-9280.1990.tb00079.x

Lee, A. Y. (2001). The mere exposure effect: An uncertainty reduction explanation revisited. Personality and Social Psychology Bulletin, 27, 1255–1266. https://doi.org/10.1177/01461672012710002

Luo, W., Feng, W., He, W., Wang, N. Y., & Luo, Y. J. (2010). Three stages of facial expression processing: ERP study with rapid serial visual presentation. Neuroimage, 49, 1857–1867. https://doi.org/10.1016/j.neuroimage.2009.09.018

Mather, M., Clewett, D., Sakaki, M., & Harley, C. W. (2016). Norepinephrine ignites local hotspots of neuronal excitation: How arousal amplifies selectivity in perception and memory. Behavioral and Brain Sciences, 39, e200. https://doi.org/10.1017/S0140525X15000667

Mende-Siedlecki, P., Said, C., & Todorov, A. (2013). The social evaluation of faces: A meta-analysis of functional neuroimaging studies. Social Cognitive and Affective Neuroscience, 8, 285–299. https://doi.org/10.1093/scan/nsr090

Mende-Siedlecki, P., Verosky, S., Turk-Browne, N. T., & Todorov, A. (2013). Robust selectivity for faces in the human amygdala in the absence of expressions. Journal of Cognitive Neuroscience, 25, 2086–2106. https://doi.org/10.1162/jocn_a_00469

Pinheiro, J., Bates, D., DebRoy, S., Sarkar, D., & R Core Team. (2017). nlme: Linear and nonlinear mixed effects models. R package version 3.1-131. Retrieved from https://cran.rproject.org/package=nlme

Rhodes, G., & Leopold, D. A. (2011). Adaptive norm-based coding of face identity. In A. J. Calder, G. Rhodes, M. H. Johnson & J. V. Haxby (Eds.), The Oxford Handbook of Face Perception (pp. 263–286).Oxford: Oxford University Press

Rhodes, G., Sumich, A., & Byatt, G. (1999). Are average facial configurations attractive only because of their symmetry? Psychological Science, 10, 52–58. https://doi.org/10.1111/1467-9280.00106

Rossion, B., Gauthier, I., Tarr, M. J., Despland, P., Bruyer, R., Linotte, S., & Crommelinck, M. (2000). The N170 occipito-temporal component is delayed and enhanced to inverted faces but not to inverted objects. NeuroReport, 11, 69–72. https://doi.org/10.1097/00001756-200001170-00014

Russell, J. A. (1980). A circumplex model of affect. Journal of Personality and Social Psychology, 39, 1161–1178. http://doi.org/10.1037/h0077714

Said, C. P., Dotsch, R., & Todorov, A. (2010). The amygdala and FFA track both social and non-social face dimensions. Neuropsychologia, 48, 3596–3605. https://doi.org/10.1016/j.neuropsychologia.2010.08.009

Said, C. P., Baron, S., & Todorov, A. (2009). Nonlinear amygdala response to face trustworthiness: Contributions of high and low spatial frequency information. Journal of Cognitive Neuroscience, 21, 519–528. https://doi.org/10.1162/jocn.2009.21041

Sánchez, Á, & Vázquez, C. (2013). Prototypicality and intensity of emotional faces using an anchor-point method. The Spanish Journal of Psychology, 16, E7. doi:10.1017/sjp.2013.9

Santos S., Almeida I., Oliveiros B., Castelo-Branco M. (2016), The role of the amygdala in facial trustworthiness processing: A systematic review and meta-analyses of fMRI studies. PLoS One, 11:e0167276. https://doi.org/10.1371/journal.pone.0167276

Satpute, A. B., Kragel, P. A., Barrett, L. F., Wager, T. D., & Bianciardi, M. (2018). Deconstructing arousal into wakeful, autonomic and affective varieties. Neuroscience Letters. https://doi.org/10.1016/J.NEULET.2018.01.042

Sergerie, K., Chochol, C., & Armony, J. L. (2008). The role of the amygdala in emotional processing: A quantitative meta-analysis of functional neuroimaging studies. Neuroscience and Biobehavioral Reviews, 32, 811–830. https://doi.org/10.1016/j.neubiorev.2007.12.002

Shen, H., Chau, D. K., Su, J., Zeng, L. L., Jiang, W., He, J., Fan, J., & Hu, D. (2016). Brain responses to facial attractiveness induced by facial proportions: Evidence from an fMRI study. Scientific Reports, 5. https://doi.org/10.1038/srep35905

Small, D. M., Gregory, M. D., Mak, Y. E., Gitelman, D., Mesulam, M. M., & Parrish, T. (2003). Dissociation of neural representation of intensity and affective valuation in human gustation. Neuron, 39, 701–711. https://doi.org/10.1016/S0896-6273(03)00467-7

Sofer, C., Dotsch, R., Wigboldus, D. H., & Todorov, A. (2015) What is typical is good the influence of face typicality on perceived trustworthiness. Psychological Science, 26, 39–47. https://doi.org/10.1177/0956797614554955

Somerville, L. H., & Whalen, P. J. (2006). Prior experience as a stimulus category confound: An example using facial expressions of emotion. Social Cognitive and Affective Neuroscience, 1, 271–274. https://doi.org/10.1093/scan/nsl040

Todorov, A. (2012). The role of the amygdala in face perception and evaluation. Motivation and Emotion, 36, 16–26. http://doi.org/10.1007/s11031-011-9238-5

Todorov, A., & Engell, A. D. (2008). The role of the amygdala in the implicit evaluation of emotionally neutral faces. Social Cognitive and Affective Neuroscience, 3, 303–312, https://doi.org/10.1093/scan/nsn033.

Tottenham, N., Tanaka, J. W., Leon, A. C., McCarry, T., Nurse, M., Hare, T. A., … Nelson, C. A. (2009). The NimStim set of facial expressions: Judgments from untrained research participants. Psychiatry Research, 168, 242–249. doi:10.1016/j.psychres.2008.05.006

Trapp, S., Schweinberger, S. R., Hayward, W. G., & Kovács, G. (2018). Integrating predictive frameworks and cognitive models of face perception. Psychonomic Bulletin & Review. https://doi.org/10.3758/s13423-018-1433-x

Valentine T. (1991). A unified account of the effects of distinctiveness, inversion, and race in face recognition. Quarterly Journal of Experimental Psychology, 43, 161–204. https://doi.org/10.1080/14640749108400966

Valentine, T., Darling, S., & Donnelly, M. (2004). Why are average faces attractive? The effect of view and averageness on the attractiveness of female faces. Psychonomic Bulletin & Review, 11, 482–487. https://doi.org/10.3758/BF03196599

Valentine, T., & Ferrara, A. (1991). Typicality in categorisation, recognition and identification: Evidence from face recognition. British Journal of Psychology, 2, 87–102.

Vuilleumier, P., Richardson, M. P., Armony, J. L., Driver, J., & Dolan, R. J. (2004). Distant influences of amygdala lesion on visual cortical activation during emotional face processing. Nature Neuroscience, 7, 1271–1278. doi:10.1038/nn1341

Willis, J., & Todorov, A. (2006). First impressions: making up your mind after a 100-ms exposure to a face. Psychological Science, 17, 592–598. https://doi.org/10.1111/j.1467-9280.2006.01750.x

Whalen, P. J., Kagan, J., Cook, R. G., Davis, F. C., Kim, H., Polis, S., … Johnstone, T. (2004). Human amygdala responsivity to masked fearful eye whites. Science, 306, 2061. https://doi.org/10.1126/science.1103617

Winston, J. S., O’Doherty, J., Kilner, J. M., Perrett, D. I., & Dolan, R. J. (2007). Brain systems for assessing facial attractiveness. Neuropsychologia, 45, 195–206. https://doi.org/10.1016/j.neuropsychologia.2006.05.009

Zald, D. H. (2003). The human amygdala and the emotional evaluation of sensory stimuli. Brain Research Reviews, 41, 88–123. https://doi.org/10.1016/S0165-0173(02)00248-5

